# Synchronous high-amplitude co-fluctuations of functional brain networks during movie-watching

**DOI:** 10.1101/2022.06.30.497603

**Authors:** Jacob C. Tanner, Joshua Faskowitz, Lisa Byrge, Daniel P. Kennedy, Olaf Sporns, Richard F. Betzel

## Abstract

Recent studies have shown that functional connectivity can be decomposed into its exact framewise contributions, revealing short-lived, infrequent, and high-amplitude time points referred to as “events.” Events contribute disproportionately to the time-averaged connectivity pattern, improve identifiability and brain-behavior associations, and differences in their expression have been linked to endogenous hormonal fluctuations and autism. Here, we explore the characteristics of events while subjects watch movies. Using two independently-acquired imaging datasets in which participants passively watched movies, we find that events synchronize across individuals and based on the level of synchronization, can be categorized into three distinct classes: those that synchronize at the boundaries between movies, those that synchronize during movies, and those that do not synchronize at all. We find that boundary events, compared to the other categories, exhibit greater amplitude, distinct co-fluctuation patterns, and temporal propagation. We show that underlying boundary events is a specific mode of co-fluctuation involving the activation of control and salience systems alongside the deactivation of visual systems. Events that synchronize during the movie, on the other hand, display a pattern of co-fluctuation that is time-locked to the movie stimulus. Finally, we found that subjects’ time-varying brain networks are most similar to one another during these synchronous events.

## INTRODUCTION

The human brain is fundamentally a complex network comprised of anatomically connected neural elements [1– 3]. This physical network constrains dynamical interactions between brain regions, inducing statistical dependencies in the activity of distant brain regions, i.e. functional connectivity (FC) [4–6]. A growing number of studies have focused on characterizing the architectural features of FC [7, 8] and linking inter-individual differences in these features to cognition [9–11], disease [12, 13], and development [14, 15].

Recent methodological advances have made it possible to precisely decompose FC into its framewise contributions [16, 17]. This “edge-centric” approach yields timevarying estimates of the co-fluctuation magnitude and valence for every pair of brain regions (edge). Previous studies have shown that, collectively, edges exhibit bursty behavior, such that long periods of quiescence are punctuated by brief, high-amplitude “events” in which many edges simultaneously and strongly co-fluctuate with one another [16, 18–24]. The whole-brain co-fluctuation patterns expressed during events are closely related to the static (time-averaged) FC [16], improve subject identification and brain-behavior correlations [16], are individualized [19], can be linked to endogenous hormone fluctuations [18], clinical status [25], and memory processes [26], and are shaped by the underlying anatomical connectivity [27].

However, the principles that determine the timing of events are undisclosed. The first “edge-centric” study showed that the whole-brain co-fluctuation amplitude is correlated during movie-watching but not at rest [16], suggesting that audiovisual stimuli may induce correlations in the timing of events. However, the spatiotemporal structure of events in naturalistic stimuli was not investigated further.

Here, we characterize this synchronization in the timing of events in greater detail with the aim of contributing a deeper understanding of the potential drivers of events. As in [16], we find that the timing of events synchronizes across participants watching the same movie. We replicate this finding using two independently-acquired, -processed, and -parcellated datasets, representing eight separate movies (each movie consisting of a small number of movie scenes, or trailers). We analyze events further, grouping them into three distinct categories: events that synchronize at the boundaries between movies, those that synchronize within the body of a movie, and those that are asynchronous. We focus further on boundary events, showing that they exhibit distinct overall co-fluctuation amplitudes, co-fluctuation pattern, and temporal structure. We show that boundary events are also underpinned by a distinct pattern of activity that involves visual processing, attentional, and control systems. Events that synchronize during movies, however, are characterized by co-fluctuation patterns that are time-locked to the movie stimulus. Additionally, we found that subjects’ time-varying brain networks are most similar to one another during these synchronous events.

Taken together, these results suggest a potential driver of one class of events, and offer a more in depth analysis of the others. Furthermore, our results suggest that the synchronization of events is more broadly related to the phenomenon of synchronization in subjects’ brains at two different scales: the whole-brain co-fluctuation amplitude, and the edge-wise pattern of co-fluctuation. Overall, our work opens up the potential for future studies to identify the psychological and biological origins of this phenomenon of event synchronization in brains.

## RESULTS

Previous studies have identified brief, high-amplitude co-fluctuations in resting-state fMRI [16, 27, 28]. The origins of these “events” are unclear. In this section, we explore the characteristics of these events in more detail using naturalistic stimuli (movie-watching) data from the Human Connectome Project [29] and a second, independently-acquired dataset [30]. In the following sections, we describe results of these analyses. Specifically, we show that events synchronize across subjects at certain parts of the movie. We provide a tripartite classification scheme for the events that occur while subjects watch movies, as follows. Synchronous boundary events occur at the end of movie scenes (and beginning of rest blocks). Synchronous movie events occur during the movie, and asynchronous events do not synchronize across subjects. We show that boundary events exhibit reproducible and distinct spatiotemporal characteristics, as well as a unique activation pattern, and finally we observe a strong positive relationship between the similarity of time-locked co-fluctuation patterns and the propensity for those time-locked frames to involve synchronous events.

### Events occur at the end of movie scenes

We first aimed to determine whether events synchronize during movie watching and, if so, when in the movies these synchronous events occurred. To investigate this, we estimated edge time series for every subject in every movie, calculated the co-fluctuation amplitude (root sum square of all edges’ at every time point; RSS), and used a previously-described algorithm to detect event frames whose RSS exceeded that of a null distribution [28]. This resulted in an “event time series” (Fig. 1a). Two of the authors (JT and RB) without reference to event time series, then manually coded and hemodynamically convolved a time series to indicate the frames at which each movie scene ended and rest blocks began (Fig. 1b; see Methods for details on the structure of this naturalistic data, and our coding protocol). Authors were blinded to brain data during coding procedure. We then “stacked” event time series and discovered an alignment between frames when many subjects had events and the indices of movie scene endings (Fig. 1c).

**FIG. 1.**
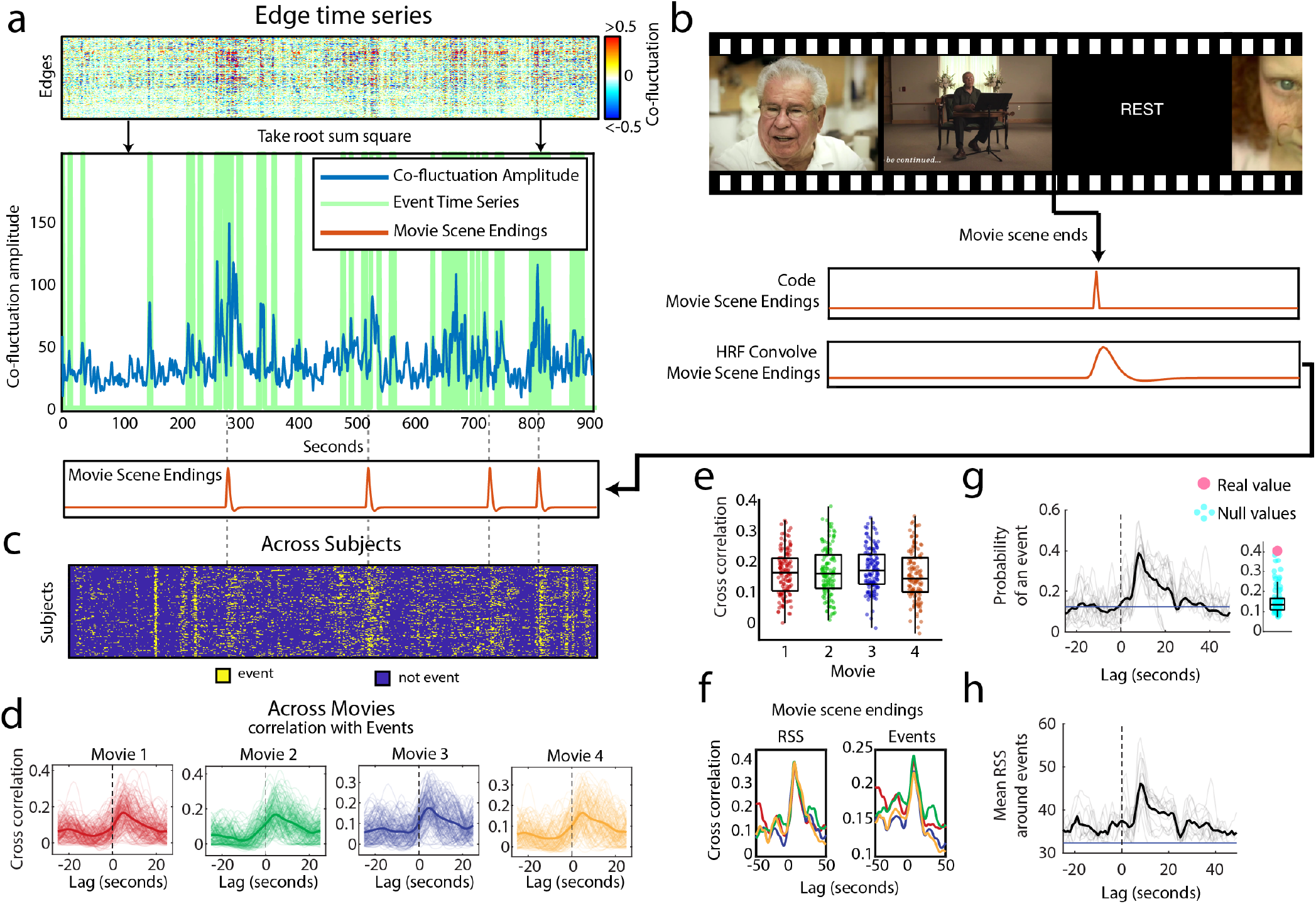
Events occurs at the end of movie scenes. (*a*) We compute edge time series (top), take the root sum square (RSS) to derive the co-fluctuation amplitude time series. Some of the peaks in co-fluctuation amplitude are determined to be events using a previously published statistical method [28]. The green line represents the binary event time series created from this method. (*b*) Schematic showing that movie scene endings were manually coded by two individuals authors (JT and RB), and then convolved with the hemodynamic response function (HRF). Example of HRF-convolved movie scene endings (orange) plotted below the relevant co-fluctuation amplitude time series. Note how many of the events from the event time series line up with movie scene endings. (*c*) Here, we “stack” the event time series across all subjects in a representative scan to illustrate that subjects tend to have events at the same moment in time, and that the moments when subjects synchronize their events often correspond to movie scene endings. This result holds across choice of FDR. (*d*) Here, we show this alignment statistically by correlating each subject’s event time series with the HRF-convolved movie scene endings while circularly shifting (referred to in figure as “Lag”) the movie scene endings in either direction. We plot this separately for each of four movies. The bold lines represent the mean across movie scene endings. (*e*) Boxplots showing the distribution of cross correlation values for the same time lag. (*f*) Plot showing the time-lagged correlation between the indices of movie scene endings, and mean RSS across subjects (first plot) per scan. Plot showing the time-lagged correlation between the indices of movie scene endings, and the number of events per time point (second plot), with one line for each movie. (*g*) Plot showing the probability of an event occurring near a movie scene ending (movie scene endings centered on zero). Black line is the probability over all scans, and the grey lines are the probability per scan. (*h*) Plot showing the mean co-fluctuation amplitude (RSS) near movie scene endings. Black line is the mean across all scans, and the grey lines are the mean per scan.

To confirm this observation statistically, we then performed several complementary tests. First, we correlated this “event time series” with the time series of hemodynamically convolved indices of movie scene endings (Fig. 1d). We found that event time series were correlated with the movie-scene endings (Fig. 1e; mean correlation per movie *r* = 0.17±0.07, *r* = 0.18±0.08, *r* = 0.18±0.07, *r* = 0.16±0.08 respectively). When we averaged event time series across subjects to obtain a group-level and pseudo-continuous estimate of event time series, we found that the fraction of subjects exhibiting an event at any given instance was maximally correlated with movie-scene endings at a lag of 5 seconds (Fig. 1f; Correlation at lag: *r* = 0.35, *r* = 0.39, *r* = 0.39, *r* = 0.40 respectively, all p-values *p <* 10^−15^). This effect is also evident in the raw and unthresholded co-fluctuation amplitude (Fig. 1f; Correlation with mean RSS across subjects at lag: *r* = 0.40, *r* = 0.45, *r* = 0.45, *r* = 0.48 respectively, all p-values *p <* 10^−15^). Finally, we found that these effects resolved to an increased probability of having an event at a lag of 5 seconds from movie-scene endings (Fig. 1g; probability of having an event *Pr*(*E*) = 0.41; compared with probability at other nearby time points; two-sample *t* -test *p <* 10^−15^).

Importantly, this result was replicated in an independently collected, processed, and parcellated data set. Instead of movie scenes, this data set presented participants with movie trailers. JT and RB again manually coded and hemodynamically convolved movie trailer endings, and compared these with the stacked event time series (Fig. S1a). We then computed the number of events within a window of 10 seconds on either side of each movie trailer ending and compared this with a null model where this window was circularly shifted 100 times. We found that there were more events near movie trailer endings than should be expected by chance (Fig. S1b,c; *p*values for each movie: *p* = 4.40×10^−3^, *p* = 1.25×10^−5^, *p* = 2.30×10^−2^, *p* = 1.26×10^−2^). Note that the 10 second window was selected to include the peak lag of 5 seconds, but also be broad enough to account for interindividual differences in timing.

Taken together, these results suggest that the ending of movies scenes (or trailers) correspond to time points when events are likely to occur in many subjects.

### Events synchronize within movie scenes

In the previous section, we demonstrated that the end of movie scenes coincide with high-amplitude cofluctuations and are often categorized as events. It is unclear, however, if synchronized co-fluctuations also occur *within* individual movies. Here, we develop a statistical test to determine whether there are other points in time, specifically during the movies, when events occur coincidentally across subjects

To address this question, we calculated the group-averaged event time series by summing across participant-level event time series. Each frame of this group time series indicated the number of subjects exhibiting an event at that instant. To identify temporally coincident events, we compared the observed group-averaged event time series with a null distribution estimated using resting-state scans rather than moviewatching (Fig. 2a). In the resting data, because subjects’ fMRI BOLD timecourses are not temporally locked to a movie stimulus, any observed synchronization is due to chance fluctuations. We also created a null model by circularly shifting event time series while subjects watched movies in order to maintain event number and relative timing while breaking alignment with the movie stimulus, but we ultimately chose the rest null model after determining it was more conservative (Fig. S7; two-sample *t* -test *p <* 10^−15^).

**FIG. 2.**
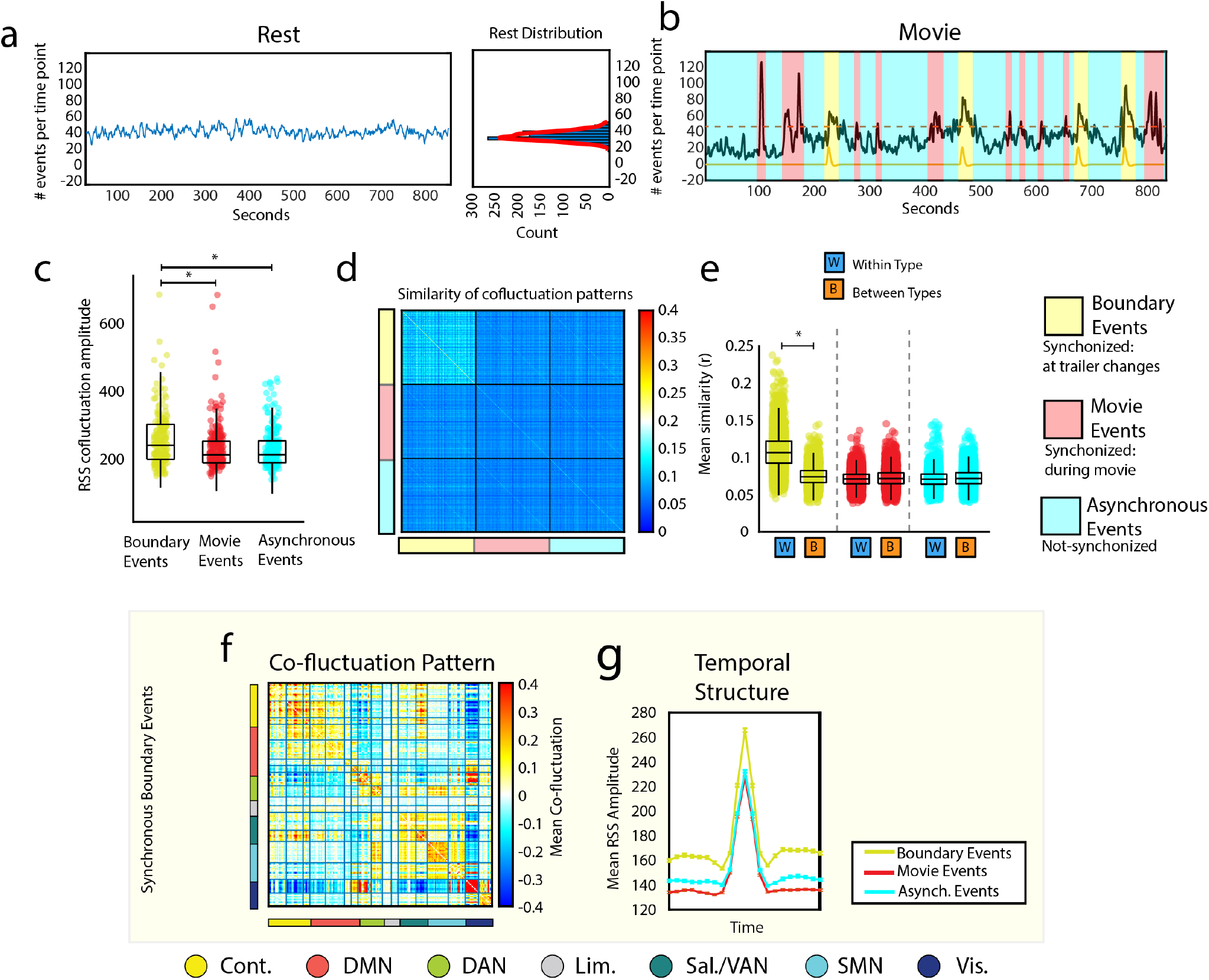
Boundary events exhibit distinct spatiotemporal characteristics. In the main text we described methods for detecting events and partitioning frames into categories based on when they occurred during the movie and the number of subjects exhibiting temporally coincident events. To detect coincident (synchronous) events, we compared event counts at each movie frame with a null distribution estimated from resting-state data. (*a*) Resting-state event count distribution (averaged across time) and event counts per frame across time. (*b*) Event counts for movie data, with frames labeled according to whether events were synchronous and occurred at movie boundaries (boundary event) or within a movie (movie event) or were not synchronized across individuals (asynchronous). Note that given the size of this cohort (*n* = 129), at least one participant exhibited an event at every frame. In principle, there could exist frames where no participant exhibited an event, necessitating a fourth category of time point (non-events). (*c*) Co-fluctuation amplitudes grouped by event type. (*d & e*) Global similarity of whole-brain co-fluctuation patterns grouped by event type (within each type, subject order is identical; mean across scans; see methods for details). (*f*) Mean co-fluctuation pattern for boundary events. (*g*) Typical temporal profile (co-fluctuation amplitude) for each event type (all events locked to their respective peaks).

Comparing the observed event time series with the null distribution of subjects at rest, we found evidence of synchronous events occurring, as expected, at movie boundaries (we refer to these as “synchronous boundary events” or simply “boundary events”; but also within movie scenes (we refer to these as “synchronous movie events” or “movie events”(we controlled for multiple comparisons by fixing the false discovery rate to *q* = 0.05, *p*_*adj*_ = 0.03). We also defined a third category of time point – those for which the number of observed events across subjects was consistent with or less than that of the null distribution (referred to as “asynchronous events”; Fig. 2b).

### Boundary events exhibit reproducible and distinct spatiotemporal characteristics

Given this tripartite classification scheme, we asked whether the three event types exhibited distinguishable characteristics. First, we tested whether boundary events exhibited dissimilar amplitudes than the other types. We found that, on average, boundary events had greater RSS than both movie and asynchronous events (Fig. 2c; twosample *t* -test *p <* 10^−15^).

Next, we asked whether the whole-brain co-fluctuation patterns expressed during events were dissimilar across event types. To do this, we calculated the similarity (bivariate linear correlation) between all pairs of detected events. Then, we averaged these scores by subject and event type. For every pair of subjects, this yielded three similarity scores – one per each event type (Fig. 2d,e). For a given event type (indicated by the blocks shown Fig. 2d), we compared within- and between-event type similarity. Intuitively, this corresponds to comparing elements in the diagonal blocks with the off-diagonal blocks. We found that “boundary events” were more similar to other boundary events than to other event types (Fig. 2e; two-sample *t* -test *p <* 10^−15^), suggesting that boundary events represent a distinct category of events, dissimilar from the other two, which are largely indistinguishable from one another.

Given that boundary events were found to be similar to one another, we took the mean across boundary events to approximate the general edge-wise co-fluctuation pattern during boundary events (Fig. 2f). Notably, boundary events displayed stronger co-fluctuation within many systems, specifically interactions of the central visual system with salience and control networks (spin test, false discovery rate fixed at *q* = 0.05, *p*_*adj*_ = 1.73 × 10^−4^; Fig. S3g).

We then examined the local temporal structure around events. Briefly, this involved temporally aligning every instance of all event types to their respective peaks (Fig. 2g). We found, in agreement with the previous RSS analysis, that at their peaks, event types were stratified based on their amplitudes, with boundary events exhibiting greater RSS than the other event types. Interestingly, we also found off-peak effects. That is, the frames before and after peaks, exhibited a similar relationship, suggesting that the frames immediately before and after boundary events also exhibited greater RSS than other event types (searched 10 frames on either side of event peak, two-sample *t* -test Bonferroni corrected *p*_*adj*_ = 4.76×10^−5^ for all 21 frames.).

Critically, all of these effects were directly replicated in a second dataset (Fig. S2). In addition, we found that the mean co-fluctuation pattern for each event type also replicated. We also found that time-averaged FC is mostly driven by synchronous movie events in the Human Connectome Project data, and that this effect is likely due to synchronous movie events containing more frames per scan than other event types (Fig. S6; two-sample *t* -test *p <* 10^−15^) but this result did not replicate in the Indiana University data set (Fig. S6e-g).

We also found evidence that the different event types contained different amounts of individualized information. Specifically, we found that in aggregate asynchronous events contained the most individualized information (Fig. S4a paired-sample *t* -test *p <* 3.37×10^−13^). On the other hand, when we considered individual frames we found that boundary events carry the most individualized information (Fig. S5; two-sample *t* -test *p <* 10^−15^; see Fig. S4 & S5 for more information).

Collectively, these results suggest that timing of events and their synchronicity across individuals shapes their amplitude, co-fluctuation pattern, identifiability and temporal evolution.

### Activation patterns during boundary events

To this point, we have analyzed co-fluctuations, which are defined at the level of edges (pairs of brain regions). While the calculation of edge-time series from activations is straightforward, obtaining the activation pattern underpinning a given co-fluctuation matrix is not. This is because every co-fluctuation matrix could have been generated by two different patterns of activity that differ in their sign at every node. For example, the product of two positively-valued nodal activations yields a positive cofluctuation. However, had the activation amplitude been identical but negatively-valued (a deactivation) we would still obtain an identical co-fluctuation. Here, we investigate the activation patterns that underlie co-fluctuations during events, focusing specifically on their configuration at movie boundaries.

First, we identified brain regions whose activity significantly increased or decreased during boundary events compared to all other time points (Fig. 3a,b; as many as 227 scans significant per region, Bonferroni corrected for number of scans tested *p*_*adj*_ = 8.77×10^−5^). Interestingly, we found that these correlations were highly system-specific, with increased activation among control b and salience/ventral attention nodes b and decreased activation in central visual (spin test; Bonferroni corrected *p*_*adj*_ = 2.90×10^−3^; Fig 3b & Fig S8). We also expanded our activation analysis to include subcortical and cerebellar regions of interest [31–34]. We found decreased activation in thalamic regions that are associated, based on their functional connectivity, with cortical control networks (Fig, 3c,e). We also found changes in activation patterns among cerebellar regions, specifically those associated with default mode, control, and dorsal attention networks (Fig. 3c,e; as many as 151 time series significant per region; Bonferroni corrected for number of scans tested *p*_*adj*_ = 8.77×10^−5^).

**FIG. 3.**
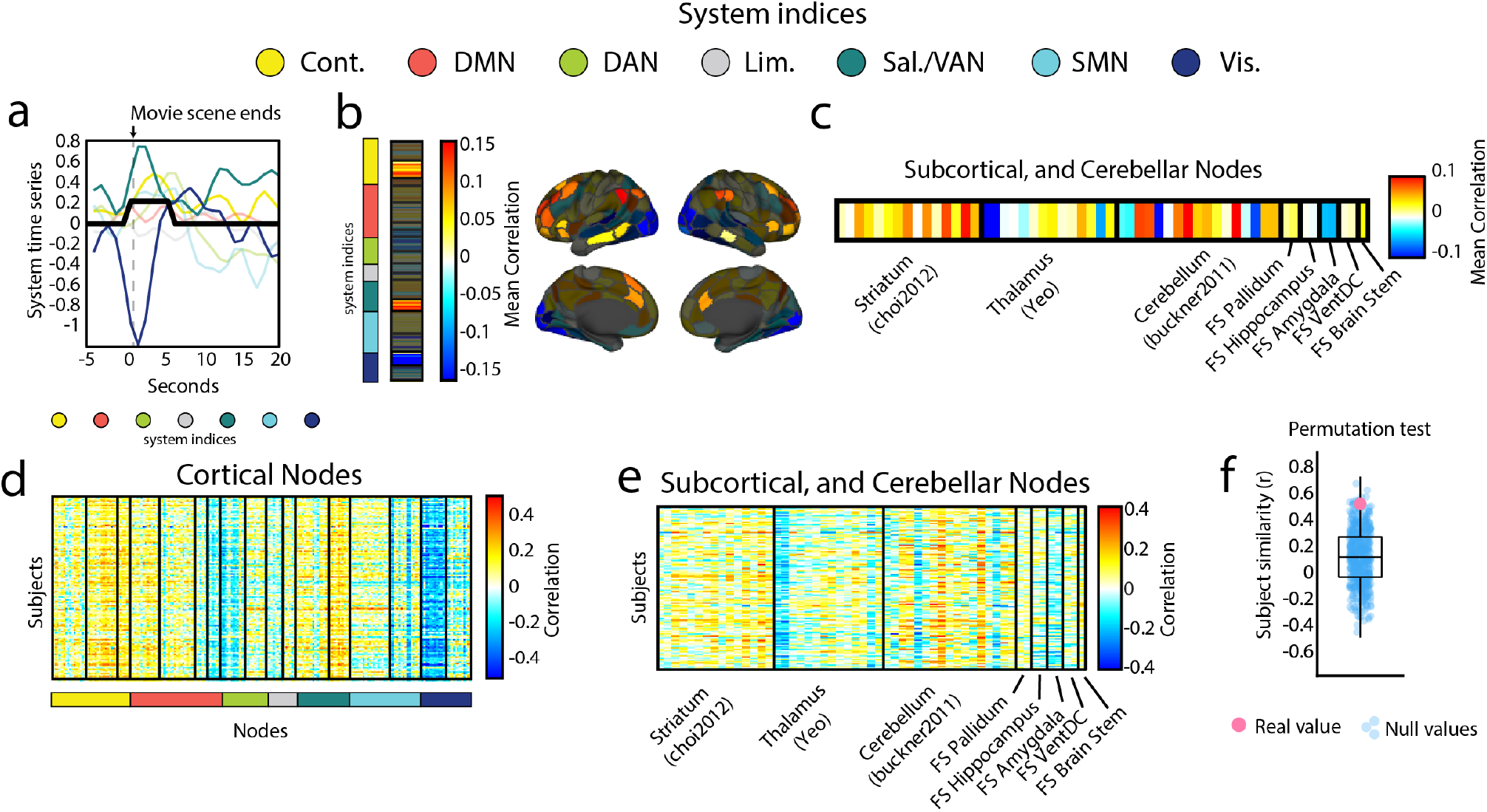
Boundary events correspond to activation of control, and salience systems, and deactivation of visual systems. (*a*) Plot of mean system activity for a window of time. The black line represents an index of boundary events. Notice the decrease in visual systems activation and an increase in salience and control. (*b*) Mean correlation values (across subjects) per region organized according to brain system (indices of brain system are color coded on the left of the plot). A space-preserving spin test was used to test if any of these brain systems tended to have a higher or lower concentration of correlation values than expected by chance. Control b, salience b, and central visual systems passed this test (*p <* 2.90 × 10^*−*3^). Systems that did not pass this test are shown shaded in grey. (*c*) Mean correlation values (across subjects) per subcortical and cerebellar region. Regions are organized mainly into parcellations for striatum, thalamus, and cerebellum. (*d & e*) Plot of the correlation values per subject and node (cortical, and subcortical respectively). (*f*) The pattern of correlation values across nodes is significantly similar across subjects, when compared with a time-shifted null model where we circularly shifted the boundary event indices before correlating them with the nodal time series (*p* = 0.03).

Importantly, we found that these correlation patterns were highly stable across individuals, both at the level of cortex as well as subcortex and cerebellum (Fig. 3d,e; mean correlation *r* = 0.51 ± 0.18, compared against correlation patterns found when using randomly permuted boundary event indices; *p <* 0.05).

In summary, these results suggest that, although cofluctuation patterns could, in principle, arise from a degenerate set of activity patterns, in the case of boundary events, they are underpinned by a single mode of activation and deactivation, involving a specific constellation of brain systems and regions.

### Time-locked movie events exhibit distinct co-fluctuation patterns

To this point we have shown that movie boundaries tend to elicit high-amplitude co-fluctuations that are synchronous across individuals and exhibit distinct spatiotemporal characteristics that distinguish them from other types of events. On one hand, perhaps this is to be expected; movie boundaries correspond to periods during the movie with similar audiovisual features, e.g. black or darkened screen and the absence of auditory stimuli. However, we have largely overlooked synchronous events that occur within the context of a single movie segment– the so-called “synchronous movie events.” In this section, we take advantage of the repeated presentation of the same movie scene across multiple scans to characterize the stable properties of synchronous and time-locked events that occur within the same movie.

To address this question, we first extracted frames corresponding to the same movie scene presented across four independent scans (a total of 516 viewings; Fig. 4a). Using the previously-described algorithm for detecting synchronous movie events, we identified and focused on three peaks (frames) in the event time series (Fig. 4b). These peaks also corresponded to points in time when subjects’ co-fluctuation patterns were highly similar (Fig. 4k) and many individuals who watched the movie scene four times reliably exhibited events upon each viewing (Fig. 4c).

**FIG. 4.**
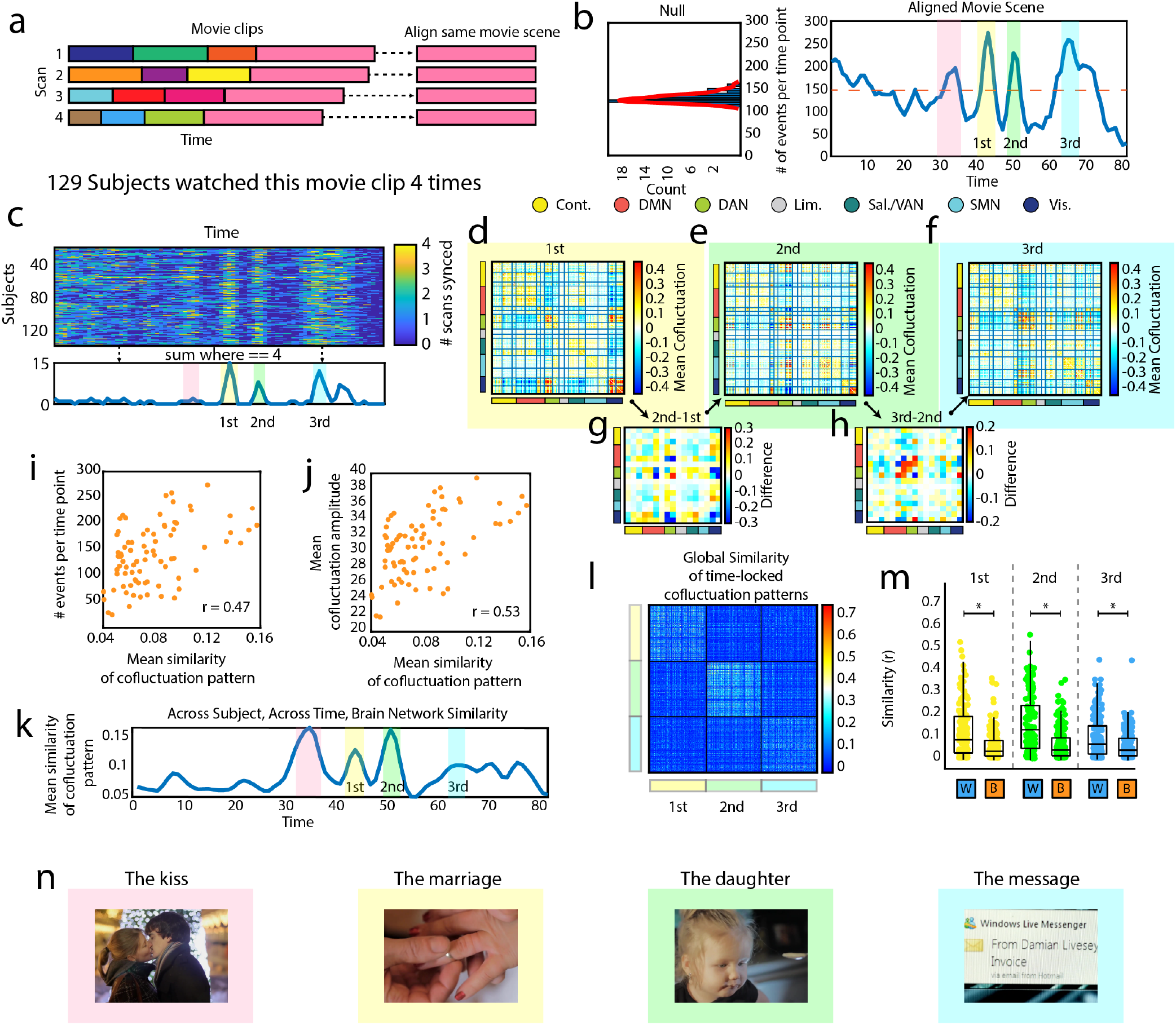
Time-locked movie events have similar co-fluctuation patterns. (*a*) Schematic showing that all four scans ended with the same movie clip, and how we temporally aligned the time series data from the four different presentations of this clip. (*b*) Histogram on the left shows the null distribution of subjects at rest. The plot on the right shows the number of events per time point in blue, with the dashed line indicating when this number is significantly greater than chance. We highlighted and numbered three peaks that we will focus on in further analyses. (*c*) This plot shows the 129 subjects who watched this movie scene four times. The values in this matrix show how many times a given subject had an event during each frame, where four is the maximum number possible. Three areas stand out in this matrix, and as the plot below the matrix shows, these areas correspond to frames where as many as 15 subjects had an event every time they watched this movie scene. (*d,e & f*) These three matrices show the mean co-fluctuation patterns (across subjects) at the frames highlighted in other plots. (*g*) This matrix shows the main differences between the 1st peak and the 2nd peak in their system by system interactions. (*h*) This matrix shows the main differences between the 2nd peak and the 3rd peak in their system by system interactions. (*i*) Plot showing the positive linear relationship between the number of events per time point, and the mean similarity of co-fluctuation patterns (across subjects). (*j*) Plot showing the positive linear relationship between the mean co-fluctuation amplitude (across subjects), and the mean similarity of co-fluctuation patterns (across subjects). (*k*) Mean similarity of co-fluctuation patterns (across subjects) plotted across time. Three frames/peaks are highlighted. (*l*) Matrix showing the similarity of co-fluctuation patterns for the three time-locked frames. (*m*) Boxplots showing the data in the previous figure divided into similarity within and between the highlighted peaks (1st, 2nd, & 3rd). Each peak is significantly more similar to itself than to the other peaks.(*n*) Screen shots from the movie scenes color coded to correspond with the time series displayed in *b,c & k*. See supplemental movie for a video of brain networks, events, and this movie side-by-side.

Next, we examined the co-fluctuation patterns during these three peaks (Fig. 4d,e,f). Interestingly, we found that these co-fluctuation patterns were dissimilar to one another, driven by system-specific differences involving the dorsal attention and temporo-parietal networks (Fig. 4g,h). This pattern is evident across subjects (Fig. 4l), where co-fluctuation patterns could be reliably distinguished from one another based on which peak they corresponded to (Fig. 4l,m; *p <* 10^−15^).

Upon reflection, this pattern – wherein peaks were dissociable from one another based on co-fluctuation patterns and event status – represents the tail of a more general phenomenon in which inter-subject similarity is positively associated with both event times and co-fluctuation amplitude (*r* = 0.47, *p* = 8.81 × 10^−6^; *r* = 0.53, *p* = 2.64 × 10^−7^, respectively; Fig. 4i,j).

In summary, these results posit a link between the similarity of co-fluctuation patterns across subjects and the propensity for those patterns to occur synchronously across individuals. Further, it reveals repeatable, finescale, and spatiotemporally dissociable event structure within movie scenes.

## DISCUSSION

Previous studies have shown that spontaneous restingstate fMRI data exhibited “events” – brief, high-amplitude, and network-level co-fluctuations. The drivers of events remain unclear. Here, we investigate this question using data acquired while subjects passively viewed movie scenes (or trailers) in the scanner. We find evidence for three types of events: boundary events, movie events, and asynchronous events. We show that boundary events are likely to occur across subjects during the boundaries between sequential movie scenes (or trailers). Further, we show that these boundary events are of greater amplitude than non-boundary events, exhibit similar co-fluctuation patterns across subjects, and follow distinct temporal trajectories. Interestingly, boundary events also carry the most subject-specific information of any event type. At the nodal level, boundary events are underpinned by a distinct pattern of activity, corresponding to deactivation of visual areas and the activation of control and ventral attention networks. In addition to showing that boundary events are distinct from other event types in many of their features, we show that movie events (synchronous events that occur during the movie) share features that are time-locked to the movies that subjects are watching. That is, when a movie scene elicits an event in many subjects, the whole-brain co-fluctuation patterns of these subjects tend to be similar. Additionally, we found that this specific pattern is a feature of a more general phenomenon wherein inter-subject synchrony in events is associated with higher inter-subject similarity in co-fluctuation patterns.

### Movie boundaries reliably elicit high-amplitude co-fluctuations

Events explain a large fraction of variance in static FC and can improve subject identification and brain-behavior correlations [16, 35]. They are individualized [28], are correlated with quotidian variability in hormones across the menstrual cycle [18], can be used to distinguish healthy controls from individuals with autism [25], and can arise spontaneously in networks whose underlying anatomical structure is modular [27]. However, the timing of events – why they occur when they do – is not understood. Resting-state datasets–the focus of most previous edge time series papers–are poorly suited for addressing this question; the unconstrained nature of rest and the absence of any temporal correlations across participants makes it difficult to establish possible event drivers.

Here, we leverage naturalistic movie-watching data [36–38] to begin addressing this question. We find evidence that, while events occur at all times during movies, they reliably and synchronously occur at the end of movie scenes. This observation is consistent with several recent studies, which link activations and intersubject synchronization at the boundaries of movies to memory [39, 40], attention processes [41], and general cognition [42].

Why events reliably occur at boundaries remains unclear. One possibility is that boundary events serve as brain-based markers for the discretization of experience [43, 44]. That is, that boundary events signify the ending of one subjectively and perceptually defined segment and the beginning of another [45–47]. Although appealing, directly testing this hypothesis necessitates the collection of additional data to confirm that movie boundaries align with participants’ subjectively defined segment boundaries [48].

Another possibility is that synchronous events arise from perturbations to a dynamical system. Many studies have shown that the temporal evolution of brain activity can be modeled as a dynamical system. If these systems are broadly similar across individuals, we might expect that driving them with the same time-varying inputs (audiovisual stimuli) will yield similar response profiles. This partially explains intersubject correlations during naturalistic paradigms. We speculate that the ends of movies, which are often typified by lower levels of luminance and sound volume, represent particularly evocative or arousing stimuli so that the shared response is brain-wide and initiates an event. Indeed, previous studies have shown that state of arousal is a powerful modulator of brain activity [49, 50] and connectivity [51, 52], suggesting that it may also play a role in event induction and should be investigated further in future studies.

We note that these hypotheses – experience segmentation and perturbations to dynamical systems – represent only two possible explanations for boundary events and are, themselves, not mutually exclusive. Additionally, we note that while movie scene boundaries are factors that explain some synchronous high-amplitude cofluctuations, we also observe synchronous events at other points within movies. The underlying mechanisms support “movie events” may be similar to boundary events, although stimuli during the movie are much richer and complex [53]. Future studies should focus on linking this second category of events with other features, e.g. movie annotations or eye-tracking data as a proxy for arousal.

### Inter-subject synchronization and events

Numerous previous studies have utilized inter-subject synchronization in brain activity to examine the roles of various brain regions in comprehending movies [53– 56]. In this research, we employ an innovative edge-centric method [16] that maintains the native temporal resolution of any time series to detect time-varying brain networks during movie-watching. This technique is known to emphasize “events,” or high-amplitude cofluctuations, which manifest as spikes in brain-wide edge amplitude. Consequently, these instances may signify substantial communication events throughout the brain.

Although these events are determined by the statistical properties of an individual subject’s brain, we discovered that all subjects’ brain networks exhibit the greatest similarity during *events* while viewing movies. In consequence, subjects not only synchronize in terms of their coarsely-defined whole-brain edge amplitude, but they also synchronize in the fluctuating strength of interregional connectivity. Importantly, this synchronized pattern of inter-regional connectivity (co-fluctuation pattern) is time-locked to the movie-stimulus suggesting that these patterns reflect brain processing of the ongoing movie. Taken together, these results suggest that the phenomenon of brain synchronization at the whole brain level during naturalistic viewing is generally connected to the occurrence of events.

### Event timing is not explained by sampling variability

Several studies have suggested that events arise not from any neurocognitively meaningful mechanism, but simply from sampling variability around a fixed correlation structure [57, 58]. That is, given a static FC matrix, nodal and edge time series must have a specific set of properties, including events. This is an important concern, and one that echoes other ongoing and unresolved debates in the network and cognitive neuroscience literature [59–61].

Our results speak directly to this controversy. Specifically, if events arise only due to stochastic fluctuations, then their occurrences should be independent and uncorrelated across individuals. However, we find that this is not the case and that, in line with previous studies [16], the timing of events within a scan session is correlated across individuals. These events cannot obviously be attributed to “rest blocks” that are interspersed between movie scenes, as we also find evidence of correlated events within movies.

These observations contribute to and enrich the ongoing dialogue surrounding exactly what features of brain networks are stationary *versus* non-stationary. Future studies – both empirical and *in silico* – should continue to investigate these and related questions.

### Future Directions

There are several ways that the results of this study could be extended in the future. For instance, we examined naturalistic stimuli (passive movie-watching). We found robust evidence of inter-subject correlations, specifically the timing of events, both during and at the end of movie scenes. While it is clear that subjects synchronize events, the precise causes remain unclear. One way to gain insight into these causes involves leveraging the movie annotations that accompany the imaging data. That is, to link the timing of events, synchronized or otherwise, to the timing of particularly salient features (e.g. objects and actions) occurring in the movie [62]. Additionally, future studies with more specified and explicit task design could also be performed to adjudicate between competing hypotheses about the origins of events. For example, online experience sampling during acquisition could be used to better understand the subjective experiences that co-occur with events [63].

Another way to extend these results would be to examine multi-modal recordings made at, potentially, faster timescales, e.g. scalp or intracranial EEG or MEG [64– 66]. Our work here, and related studies, have focused almost exclusively on fMRI data. However, fMRI has a number of known limitations, most notably it exhibits relatively poor temporal resolution, which may obscure rapidly occurring neural processes, e.g. fine-grained temporal structure that occurs just prior to or following an event [67–69]. Future studies should investigate inter-synchrony of events using imaging modalities that are acquired at faster rates.

Here, we focus on empirical recordings. However, recent studies have shown that events can be observed in synthetic data generated by dynamic models [27]. This opens up several avenues for future work, including artificially “stimulating” *in silico* brains to induce events and, if we have two independent simulations, whether simultaneous stimulation leads them to synchronize [70]. The simulation-based analyses also make it possible to investigate and test related hypotheses using data from non-human subjects, e.g. macaque [71].

One of our key findings was that events are temporally coincident during movie-watching and especially near the boundaries between movie trailers. However, participants consistently exhibit events throughout the scan, even if the timing is not coincident with other individuals. What are the origins of these “asynchronous events?” One possibility is that they reflect an intrinsic mode of rest that is decoupled from the ongoing task/movie stimuli [72]. Future studies, likely with dedicated experimentation, are needed to clarify the distinction between asynchronous and synchronous events.

Another noteworthy point concerns how events were defined. Here, a time point was considered an event if global – i.e. whole-brain – cofluctuations exceeded a statistical criterion. We found that these global events reliably occurred during movie boundaries, which correspond to large shifts in movie content. Future studies should investigate locally-defined events – e.g. at the level of brain systems – which may be linked to more subtle variation in movie features.

Finally, while we find evidence of synchronous events, we note that synchronization does not imply that all subjects experienced an event at the same moment (just that a sufficiently large number did). An important open question is whether those subjects that did not experience an event are distinguished from their peers along other dimensions as well, e.g. cognitive, clinical, demographic profiles. For example, perhaps subjects that experienced an event have better recall of related movie scenes than subjects that did not.

## Limitations

There are several limitations of the current study. One notable issue is the use of fMRI data to infer brain activity. Functional imaging data is fundamentally an indirect (and slow) measure of the hemodynamic response to population-level activity. Follow-up studies should aim to reproduce these results using more direct assessments of activity, e.g. intracranial EEG.

Additionally, there are several limitations associated with the movie data themselves. The first concerns the subjectivity with which individuals experience the movie stimuli. While the stimuli are identical across individuals, their experiences may not be, e.g. participants may attend to different movie features at the same time point or draw on their personal history, leading to dissimilar experience. Without subjective post-scan reports or incorporation of additional data modalities, e.g. eye-tracking, it remains challenging to uncover the drivers of synchronized events.

Furthermore, it is important to clarify that while previous studies have primarily focused on events during the resting-state, we focus here on events detected during passive movie-watching. While the statistical approach used to detect periods of high-amplitude co-fluctuations is identical, it remains unclear whether resting and movie events share common neurocognitive underpinnings. Future studies should investigate their differences and commonalities in more detail.

Lastly, there remain challenges related to data processing and analysis. Specifically, the procedures for disambiguating activations (first-order effects) from co-fluctuations (second-order effects) during moviewatching watching are not well defined. In blocked or event-related task design, regressors can be constructed to orthogonalize time series with respect to activations [73], making it possible to effectively remove activations as confounds prior to estimating connectivity or cofluctuations. However, with movie-watching data, stimuli are presented in a naturalistic way, meaning that many stimuli overlap and co-occur, e.g. the presence of a face is often correlated with speech, making it difficult to control for all possible stimuli. Moreover, the stimuli are presented in a continuous stream, similarly making it difficult to construct appropriate regressors. This remains an active and contentious area of research. Future methodological studies will address this potential issue. One particular concern unique to our findings is the presence of “rest blocks” in the HCP dataset. An alternative interpretation of our results is that because the transition from a “movie state” to “rest state” is accompanied by a short pause, it is the pause itself that induces a non-stationarity in the activations, yielding a boundary event.

## Conclusion

In conclusion, this study represents an analysis of the synchronization of high-amplitude co-fluctuations, or “events”, while participants are watching the same movie in two separately collected, processed and parcellated data sets. Our analysis suggests that events during movie watching can be divided into three categories based upon their synchronization. Synchronous boundary events occur at the end of movie scenes, or trailers, and are distinct from the other event types in overall co-fluctuation pattern, amplitude and temporal structure, as well as carrying the most subject-specific information at the level of individual frames. Synchronous movie events occur during the movie and their time-locked co-fluctuation patterns are highly similar across subjects. In fact, we found a general relationship between the degree of event synchronization across subjects and the across subject similarity in co-fluctuation pattern. These two types of synchronous events suggests that events can be elicited by environmental stimuli in a similar manner across subjects. Asynchronous events also occur throughout the movie, suggesting an individualized element to the inducement of events by movies. Indeed, when considered together asynchronous events also carry the most individualized information about subjects.

## MATERIALS AND METHODS

### Human Connectome Project Data

The Human Connectome Project (HCP) 7T dataset [29] consists of structural magnetic resonance imaging (T1w), resting state functional magnetic resonance imaging (rsfMRI) data, movie watching functional magnetic resonance imaging (mwfMRI) from 184 adult subjects. These subjects are a subset of a larger cohort of approximately 1200 subjects additionally scanned at 3T. Subjects’ 7T fMRI data were collected during four scan sessions over the course of two or three days at the Center for Magnetic Resonance Research at the University of Minnesota. Subjects’ 3T T1w data collected at Washington University in St. Louis. The study was approved by the Washington University Institutional Review Board and informed consent was obtained from all subjects.

### Demographics

We analyzed MRI data collected from *N*_*s*_ = 129 subjects (77 female, 52 male), after excluding subjects with poor quality data. Our exclusion criteria was as follows: where each spike is defined as relative framewise displacement of at least 0.25 mm, we excluded subjects who fulfill at least 1 of the following criteria: greater than 15% of time points spike, average framewise displacement greater than 0.2 mm; contains any spikes larger than 5mm. Following this filter, subjects who contained all four scans were retained. At the time of their first scan, the average subject age was 29.36±3.36 years, with a range from 22−36. 70 of these subjects were monozygotic twins, 57 where non-monozygotic twins, and 2 were not twins.

### MRI acquisition and processing

A comprehensive description of the imaging parameters and image preprocessing can be found in [74] and in HCP’s online documentation (https://www.humanconnectome.org/study/hcp-youngadult/document/1200-subjects-data-release). T1w were collected on a 3T Siemens Connectome Skyra scanner with a 32-channel head coil. Subjects underwent two T1-weighted structural scans, which were averaged for each subject (TR = 2400 ms, TE = 2.14 ms, flip angle = 8°, 0.7 mm isotropic voxel resolution). fMRI were collected on a 7T Siemens Magnetom scanner with a 32channel head coil. All 7T fMRI data was acquired with a gradient-echo planar imaging sequence (TR = 1000 ms, TE = 22.2 ms, flip angle = 45°, 1.6 mm isotropic voxel resolution, multi-band factor = 5, image acceleration factor = 2, partial Fourier sample = 7/8, echo spacing = 0.64 ms, bandwidth = 1924 Hz/Px). Four resting state data runs were collected, each lasting 15 minutes (frames = 900), with eyes open and instructions to fixate on a cross. Four movie watching data runs were collected, each lasting approximately 15 minutes (frames = 921, 918, 915, 901), with subjects passively viewing visual and audio presentations of movie scenes. Movies consisted of both freely available independent films covered by Creative Commons licensing and Hollywood movies prepared for analysis [75]. For both resting state and movie watching data, two runs were acquired with posterior-to-anterior phase encoding direction and two runs were acquired with anterior-to-posterior phase encoding direction.

Structural and functional images were minimally preprocessed according to the description provided in [74]. 7T fMRI images were downloaded after correction and reprocessing announced by the HCP consortium in April, 2018. Briefly, T1w images were aligned to MNI space before undergoing FreeSurfer’s (version 5.3) cortical reconstruction workflow. fMRI images were corrected for gradient distortion, susceptibility distortion, and motion, and then aligned to the corresponding T1w with one spline interpolation step. This volume was further corrected for intensity bias and normalized to a mean of 10000. This volume was then projected to the 2mm *32k fs LR* mesh, excluding outliers, and aligned to a common space using a multi-modal surface registration [76]. The resultant cifti file for each HCP subject used in this study followed the file naming pattern: * Atlas MSMAll hp2000 clean.dtseries.nii. Resting state and moving watching fMRI images were nuisance regressed in the same manner. Each minimally preprocessed fMRI was linearly detrended, band-pass filtered (0.008-0.25 Hz), confound regressed and standardized using Nilearn’s signal.clean function, which removes confounds orthogonally to the temporal filters. The confound regression strategy included six motion estimates, mean signal from a white matter, cerebrospinal fluid, and whole brain mask, derivatives of these previous nine regressors, and squares of these 18 terms. Spike regressors were not applied. Following these preprocessing operations, the mean signal was taken at each time frame for each node, as defined by the Schaefer 400 parcellation [77] in *32k fs LR* space.

### Indiana University Data Demographics

We analyzed MRI data collected from *N*_*s*_ = 29 subjects (5 female, 24 male; 25 were right-handed). This cohort was male-dominant, as subjects were intended to serve as controls for a study in autism spectrum disorder, which is more common in men than women. At the time of their first scan, the average subject age was 24.9± 4.7 years.

### MRI acquisition and processing

MRI images were acquired using a 3T whole-body MRI system (Magnetom Tim Trio, Siemens Medical Solutions, Natick, MA) with a 32-channel head receive array. Both raw and prescan-normalized images were acquired; raw images were used at all preprocessing stages and in all analyses unless specifically noted. During functional scans, T2*-weighted multiband echo planar imaging (EPI) data were acquired using the following parameters: TR/TE = 813/28 ms; 1200 vol; flip angle = 60°; 3.4 mm isotropic voxels; 42 slices acquired with interleaved order covering the whole brain; multi-band acceleration factor of 3. Preceding the first functional scan, gradient-echo EPI images were acquired in opposite phase-encoding directions (10 images each with P-A and A-P phase encoding) with identical geometry to the EPI data (TR/TE = 1175/39.2 ms, flip angle = 60°) to be used to generate a fieldmap to correct EPI distortions, similar to the approach used by the Human Connectome Project [78]. High-resolution T1-weighted images of the whole brain (MPRAGE, 0.7 mm isotropic voxel size; TR/TE/TI = 2499/2.3/1000 ms) were acquired as anatomical references.

All functional data were processed according to an in-house pipeline using FEAT (v6.00) and MELODIC (v3.14) within FSL (v. 5.0.9; FMRIB’s Software Library, www.fmrib.ox.ac.uk/fsl), Advanced Normalization Tools (ANTs; v2.1.0) [79], and Matlab R2014b. This pipeline was identical to the **GLM + MGTR** procedure described in [30].

In more detail, individual anatomical images were bias-corrected and skull-stripped using ANTs, and segmented into gray matter, white matter, and CSF partial volume estimates using FSL FAST. A midspace template was constructed using ANTs’ *buildtemplateparallel* and subsequently skull-stripped. Composite (affine and diffeomorphic) transforms warping each individual anatomical image to this midspace template, and warping the midspace template to the Montreal Neurological Institute MNI152 1mm reference template, were obtained using ANTs.

For each functional run, the first five volumes (≈4 seconds) were discarded to minimize magnetization equilibration effects. Framewise displacement traces for this raw (trimmed) data were computed using *fsl motion outliers*. Following [30, 80], we performed FIX followed by mean cortical signal regression. This procedure included rigid-body motion correction, fieldmap-based geometric distortion correction, and non-brain removal (but not slice-timing correction due to fast TR [78]). Preprocessing included weak highpass temporal filtering (*>*2000 s FWHM) to remove slow drifts [78] and no spatial smoothing. Off-resonance geometric distortions in EPI data were corrected using a fieldmap derived from two gradient-echo EPI images collected in opposite phase-encoding directions (posterior-anterior and anterior-posterior) using FSL topup.

We then used FSL-FIX [81] to regress out independent components classified as noise using a classifier trained on independent but similar data and validated on hand-classified functional runs. The residuals were regarded as “cleaned” data. Finally, we regressed out the mean cortical signal (mean BOLD signal across gray matter partial volume estimate obtained from FSL FAST). All analyses were carried out on these data, which were registered to subjects’ skull-stripped T1-weighted anatomical imaging using Boundary-Based Registration (BBR) with *epi reg* within FSL. Subjects’ functional images were then transformed to the MNI152 reference in a single step, using ANTS to apply a concatenation of the affine transformation matrix with the composite (affine + diffeomorphic) transforms between a subject’s anatomical image, the midspace template, and the MNI152 reference. Prior to network analysis, we extracted mean regional time series from regions of interest defined as sub-divisions of the 17-system parcellation reported in [82] and used previously [83–85]. Wakefulness during movie and rest scans was monitored in real-time using an eye tracking camera (Eyelink 1000).

### Naturalistic stimuli and coding boundaries

#### Human Connectome Project Data

Movies consisted of concatenated movie scenes with 20 second blocks of rest between them. The movies scenes were sourced from both freely available independent films covered by Creative Commons licensing and Hollywood movies prepared for analysis [75].

Both JCT and RFB manually coded the boundaries between the end of movie scenes and the beginning of rest blocks and confirmed consistency across codings. Additionally, JCT confirmed that these codings aligned with the end of movie scenes and the beginning of rest blocks using the RGB values from a digitized version of the movies. The movie screen during rest blocks, on average, contains more black pixels (See Fig 1b for example). By indexing the number of black pixels and overlaying the coded boundaries, JCT found an alignment between the beginning of rest blocks (where the number of black pixels increased) and coded boundaries.

#### Indiana University Data

All movies were obtained from Vimeo (https://vimeo.com). They were selected based on multiple criteria. First, to ensure that movie trailers represented novel stimuli, we excluded any movie that had a wide theatrical release. Secondly, we excluded movies with potentially objectionable content including nudity, swearing, drug use, etc. Lastly, we excluded movies with intentionally startling events that could lead to excessive in-scanner movement.

Each trailer lasted between 45 and 285 seconds (approximately 1 to 5 minutes). Each movie scan comprised between four and six trailers with genres that included documentaries, dramas, comedies, sports, mystery, and adventure (See Table. S1 for more details). Both JCT and RFB manually coded the boundaries between trailers and confirmed consistency across codings.

### Functional connectivity

Functional connectivity (FC) measures the statistical dependence between the activity of distinct neural elements. In the modeling of macroscale brain networks with fMRI data, this usually means computing the Pearson correlation of brain regions’ activity time series. To calculate FC for regions *i* and *j*, then, we first standardize their time series and represent them as zscores. We denote the z-scored time series of region *i* as **z**_*i*_ = [*z*_*i*_(1), …, *z*_*i*_(*T*)], where *T* is the number of samples. The Pearson correlation is then calculated as:

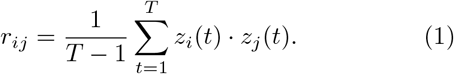

In other words, the correlation is equal to the temporal average of two regions’ co-fluctuation.

### Edge time series

We analyzed edge time series data. Edge time series can be viewed as a temporal decomposition of a correlation (functional connection) into its framewise contributions. Note that Pearson correlation is calculated as 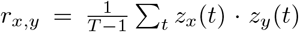, where *T* is the number of samples and 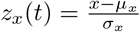 is the z-scored transformation of the time series *x* = [*x*(1), …, *x*(*T*)]. If we omit the summation in our calculation of *r*_*x,y*_, we obtain a time series *r*_*x,y*_(*t*) = *z*_*x*_(*t*) *z*_*y*_(*t*), whose elements index the instantaneous co-fluctuation between variates *x* and *y*. Here, we estimated the edge time series for all pairs of brain regions {*i, j*}.

We analyzed edge time series using two distinct approaches.

#### Edge time series amplitude (RSS)

First, we calculated the amplitude at each frame as the root sum of squared co-fluctuations. That is, the amplitude at time *t* was given by:

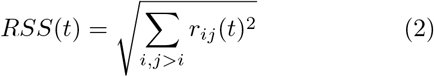

#### Whole-brain co-fluctuation patterns

The section way in which we analyzed edge time series was by considering the whole-brain co-fluctuation pattern at a given time point. That is, we focused on the cofluctuation matrices, *C*(*t*) ∈ ℝ ^*N* ×*N*^. The element *i, j* in this matrix denoted the co-fluctuation magnitude between regions *i* and *j* at time *t*, i.e. *r*_*ij*_(*t*). Although co-fluctuation matrices are not correlation matrices, they can nonetheless be analyzed using similar network science tools.

### Global Similarity Matrix

We assessed the similarity of events between subjects and types (boundary, movie, and asynchronous). The procedure for doing so involved two steps. First, for a given pair of subjects, *r* and *s*, with *n*_*r*_ and *n*_*s*_ events, respectively, we calculated the 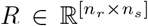 similarity matrix between all pairs of events. The elements of this matrix could be further aggregated and averaged based on the category to which events were assigned, reducing *R* to an asymmetric, [3×3] matrix (rows and columns correspond to the three event types).

This first step was repeated for all pairs of subjects, each time yielding an analogous [3×3] matrix. At its completion this, second step yielded a 387×387 similarity matrix, whose rows and columns could be reordered by event types. We show this matrix in Fig. 2d.

### Co-fluctuation Matrix Differences

In the main text, we described differences in the co-fluctuation matrices associated with different event types. To assess these differences, we performed the following set of calculations. For a given pair of event types, we identified all instances of each type and calculated, at the level of edges, the differences in their co-fluctuation magnitude and the mean sign of those differences. We then retained the sign of these differences. The result is a *n*×*n* matrix for each subject whose elements are {−1, 1}. Note that this calculation was performed within subjects.

Next, we examined whether, across subjects, edges exhibited concordant behavior using a paired-samples *t* test. That is, we assessed whether, across subjects, an edge’s co-fluctuation magnitude under one event type was consistently greater than or less than that of another event type. This procedure was repeated for every pair of event types.

## Author Contributions

JCT, OS, and RFB conceived of experiments, designed analyses, wrote the initial versions of the manuscript, and edited the manuscript. JCT performed all analyses. JF, LB, and DK contributed data. All authors helped edit the manuscript.

## Acknowledgements

Data were provided [in part] by the Human Connectome Project, WU-Minn Consortium (Principal Investigators: David Van Essen and Kamil Ugurbil; 1U54MH091657) funded by the 16 NIH Institutes and Centers that support the NIH Blueprint for Neuroscience Research; and by the McDonnell Center for Systems

Neuroscience at Washington University. This material is based on work supported by NSF Grant IIS-2023985 (RFB, OS), NSF Grant 1735095, Interdisciplinary Training in Complex Networks (JCT), and Indiana University Office of the Vice President for Research Emerging Area of Research Initiative, Learning: Brains, Machines and Children (RFB). This research was supported in part by Lilly Endowment, Inc., through its support for the Indiana University Pervasive Technology Institute.

---

**TABLE S1.** Movies included in each movie scan for the Indiana University dataset.

| Scan | Title | Genre | Runtime |
| --- | --- | --- | --- |
| 1 | Man Up and Go | documentary/emotional | 4m20s |
| 1 | The First 70 | documentary | 3m |
| 1 | Fixation | documentary/adventure | 1m42s |
| 1 | The Living | drama | 2m |
| 1 | SAMSARA | documentary/"unparalleled sensory experience" | 1m35s |
| 1 | Blood Brother | documentary | 2m20s |
| 2 | Birdmen | documentary/adventure | 3m59s |
| 2 | Groomed | drama | 1m30s |
| 2 | Cold | outdoor/sports | 2m |
| 2 | Sleepwalkers | drama | 2m |
| 2 | A Kind of Show | comedy | 1m |
| 3 | Geofish | documentary/adventure | 4m40s |
| 3 | The Debut | outdoor/sports | 3m23s |
| 3 | Dreams of a Life | documentary/mystery | 2m10s |
| 3 | The Front Man | documentary | 2m30s |
| 3 | This Is Vanity | drama | 1m |
| 4 | Planetary | documentary | 4m30s |
| 4 | Sign Painters | documentary | 2m50s |
| 4 | Florida Man | documentary/drama | 2m |
| 4 | The Sleeping Bear | drama | 3m40 |

**FIG. S1.**
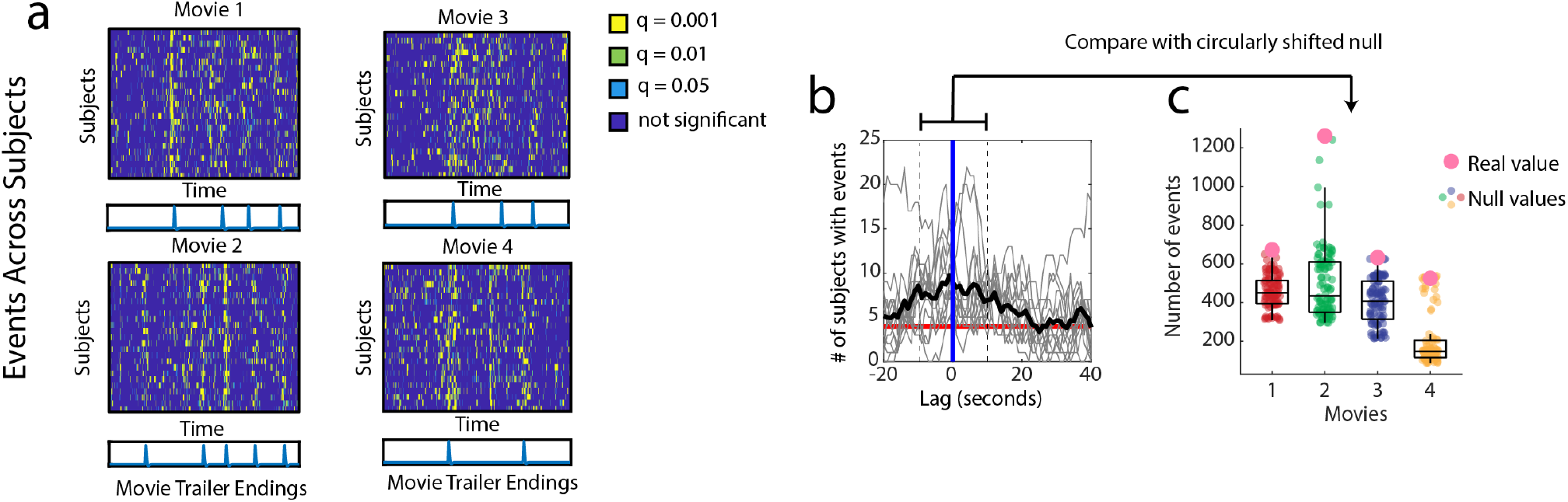
Replication of synchronous boundary events in Indiana University data set. (*a*) Each panel represents two things: 1) the stacked event time series of all subjects, and 2) the hrf-convolved movie trailer endings. Each of the four figures is representative of a different scan/movie-presentation for the Indiana University data set. (*b*) Plot showing the number of subjects with an event for each trailer ending (for every scan). Bold line indicates the mean across all trailer endings. (*c*) Using a window of 10 seconds on either side of all movie trailer endings, we compared the number of events within this window to a null model where this window is circularly shifted 100 times.

**FIG. S2.**
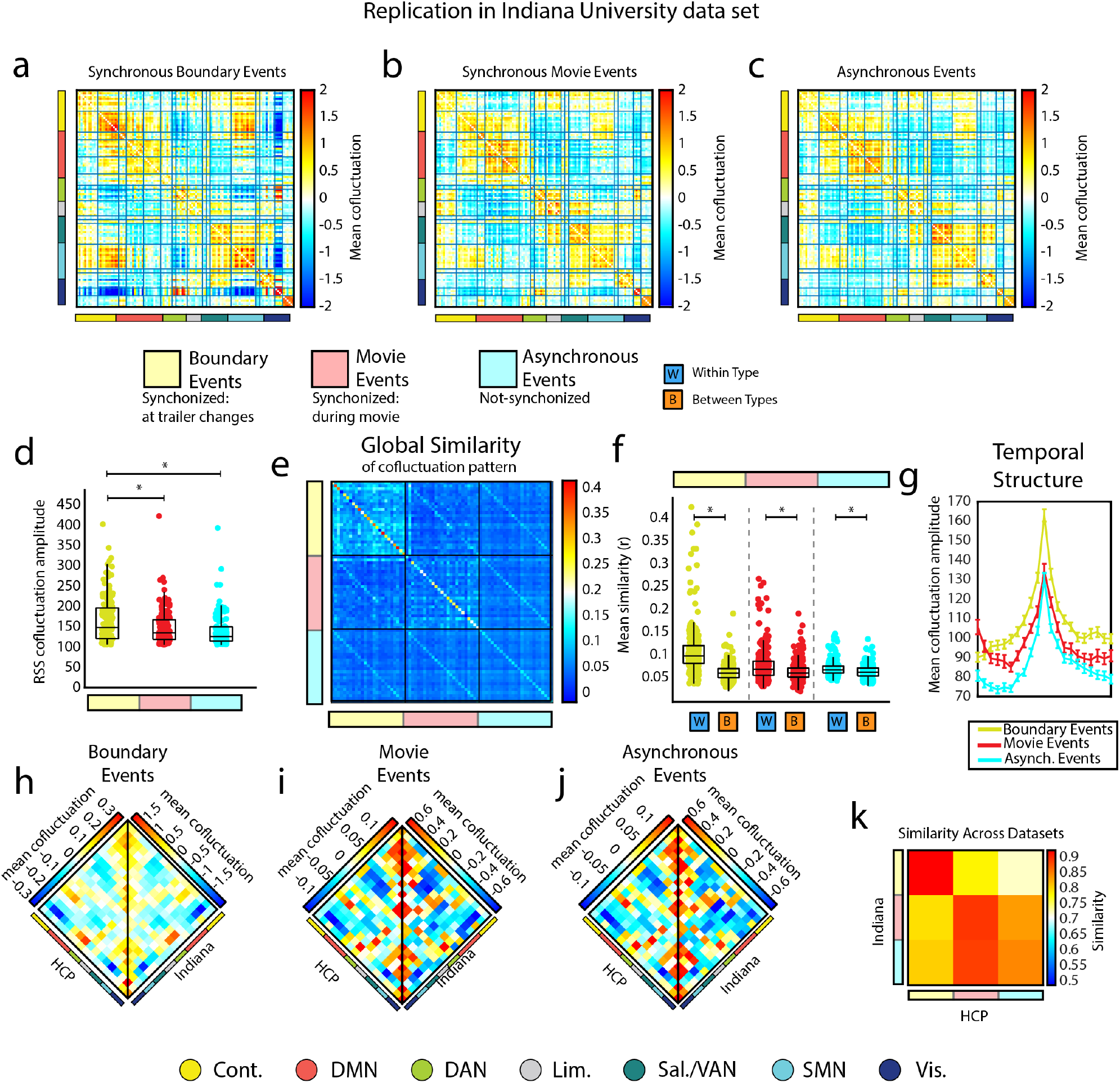
Three types of events: replication in Indiana University data set. (*a-c*) Mean co-fluctuation matrices representing the mean of all events for a given event type. (*d*) Boxplots of the distribution of co-fluctuation amplitudes per event type. (*e*) Global similarity within and between different event types (global similarity is defined in the methods section). (*f*) Boxplots showing the data from the previous figure divided by similarity values within the same type and similarity values between different types. (*g*) Plot showing the mean temporal trajectory of each event type. (*h-j*) System by system mean co-fluctuation patterns across boundary events, movie events, and asynchronous events respectively for both data sets (HCP: left; Indiana: Right). (*k*) Across data set similarity of system by system mean cofluctuation matrices for each event type.

**FIG. S3.**
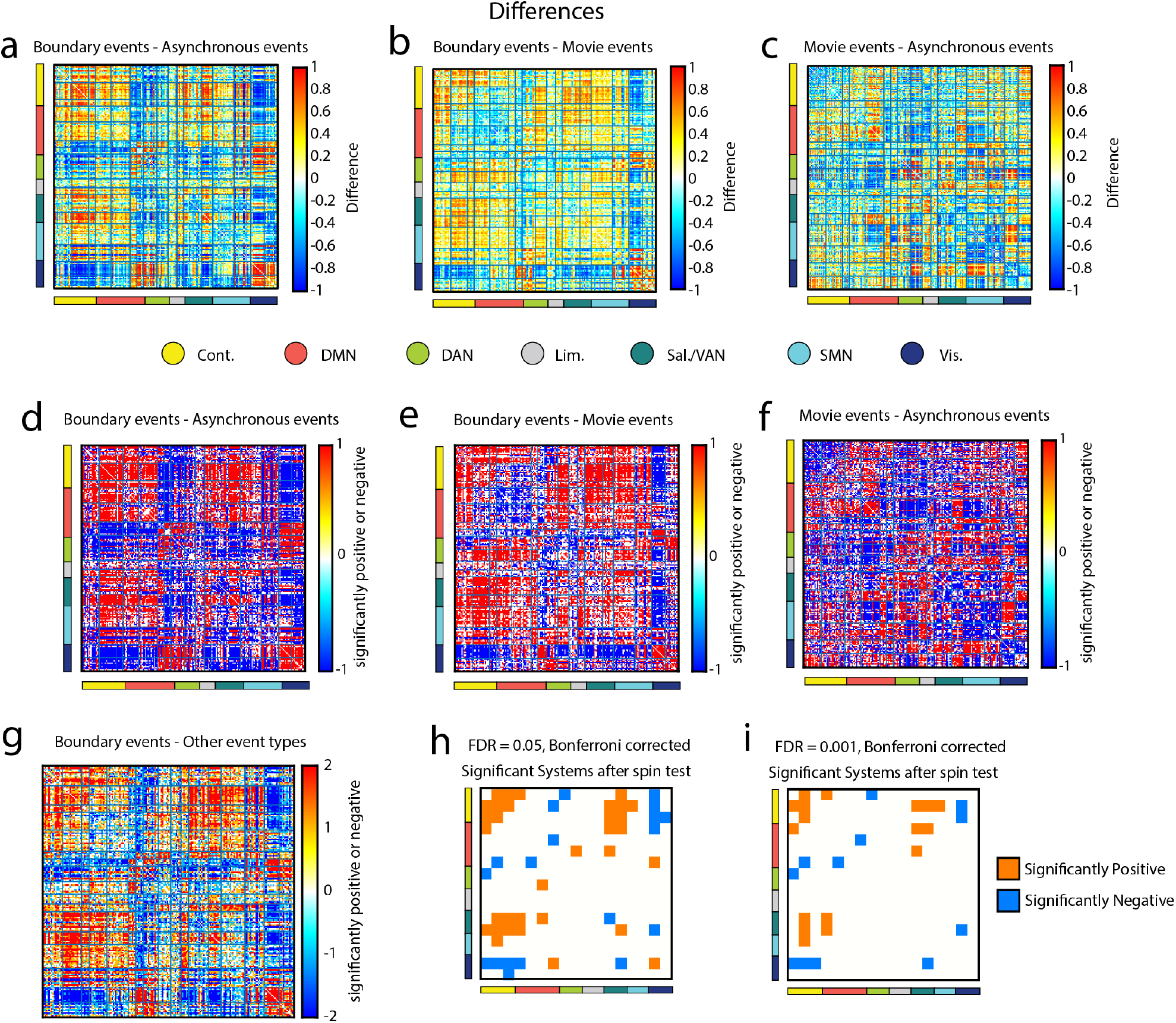
Differences between the Event Types. All differences described here are computed using the method detailed in the methods section. (*a*) Difference between boundary events and asynchronous events. (*b*) Difference between boundary events and movie events. (*c*) Difference between movie events and asynchronous events. (*d*) Significantly different edges between boundary events and asynchronous events each colored according to whether the difference was significantly positive or significantly negative. (*e*) Significantly different edges between boundary events and movie events each colored according to whether the difference was significantly positive or significantly negative. (*f*) Significantly different edges between movie events and asynchronous events each colored according to whether the difference was significantly positive or significantly negative. (*g*) Significantly different edges between boundary events and both asynchronous events as well as movie events each colored according to whether the difference was significantly positive or significantly negative. (*h*) Significant system by system edges after running a space-preserving null model (spin test) to test if the significant edges in the previous figure are more concentrated in system by system blocks than we should expect by chance. Here we controlled for multiple comparisons by fixing the false discovery rate to *q* = 0.05. (*i*) Same test as previous figure with a false discovery rate set to *q* = 0.001.

**FIG. S4.**
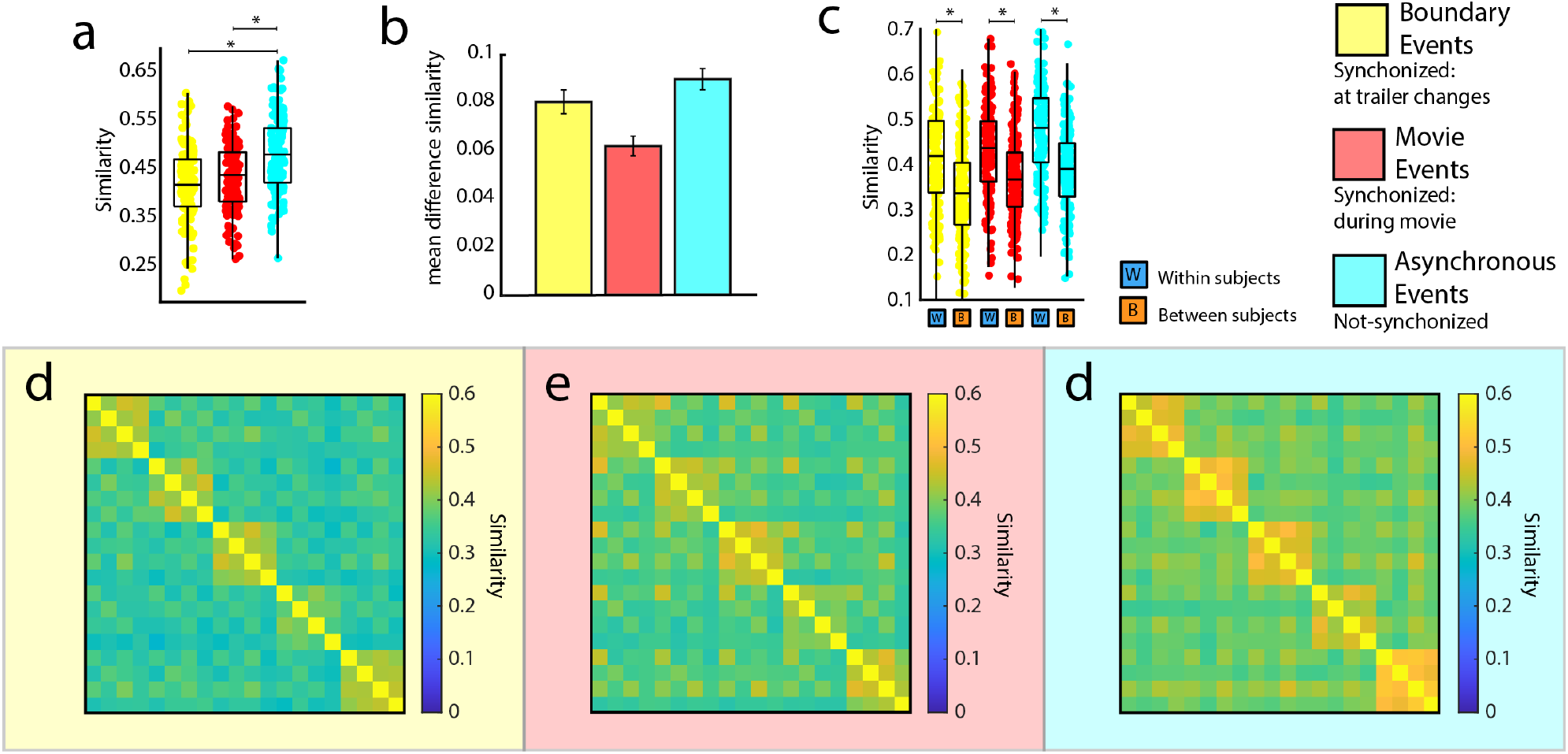
Mean co-fluctuation patterns of synchronous events carry more subject-specific information than other event types. Following research that suggests that events offer increased identifiability in human brain networks[19], we explored the identifiability of each event type. In these analyses, we derived mean co-fluctuation patterns for each type of event and for each scan. The different event types occurred at different rates. To ensure that mean co-fluctuation patterns were estimated using similar number of samples, we sub-sampled from the total number of frames assigned to each event type and calculated the mean co-fluctuation pattern using only those frames. (*a*) We find that asynchronous events were more similar within subjects compared to boundary or movie events (paired-sample *t* -test *p <* 3.37*×*10^*−*13^). While all event types maintain significant individualized features such that within subject similarity is higher than between subject similarity (*c*; two-sample *t* -test *p <* 10^*−*15^), we find that asynchronous events maintain the greatest difference in within versus between-subject similarity (*b*) [Barplots showing the difference between the means of the within-subject similarity distribution and the mean of the between-subject similarity distribution for each event type. Error bars represent the standard error of the difference between the two means.] Panels (*d-f*) show similarity matrices representing a set of 5 subjects for each event type. Blocks along the diagonal indicate within-subject similarity across 4 scans. Off-diagonal elements indicate between-subject similarity. In order to display most of the subjects while still maintaining the visual intuition that these plots provide with 5 subjects, we took the mean of these matrices for 25 non-overlapping windows into the full similarity matrix (representing a total of 125 out of the 129 subjects). Notice how the within-subject blocks are strongest in asynchronous events. We also found strong subject-level effects across event types when events are considered individually instead of taking the mean (Fig. S5; synchronous movie events did not pass identifiability tests for two scans). To confirm this, we directly compared the within-subject similarity to between-subject similarity for every frame of every event type separately (two-sample *t* -test *p <* 8.66 *×* 10^*−*4^). Interestingly, we found that – when considered individually – identifiability was actually greater in boundary events (Fig. S5; two-sample *t* -test *p <* 10^*−*15^). This difference in the relative identifiability of event types is likely due to the overall similarity of boundary events to one another (recall Fig. 2d,e). That is, the pattern of co-fluctuation found in asynchronous events is more diverse, but in aggregate asynchronous events pull together more identifiable information, while on an individual basis more identifiable information can be found in boundary events.

**FIG. S5.**
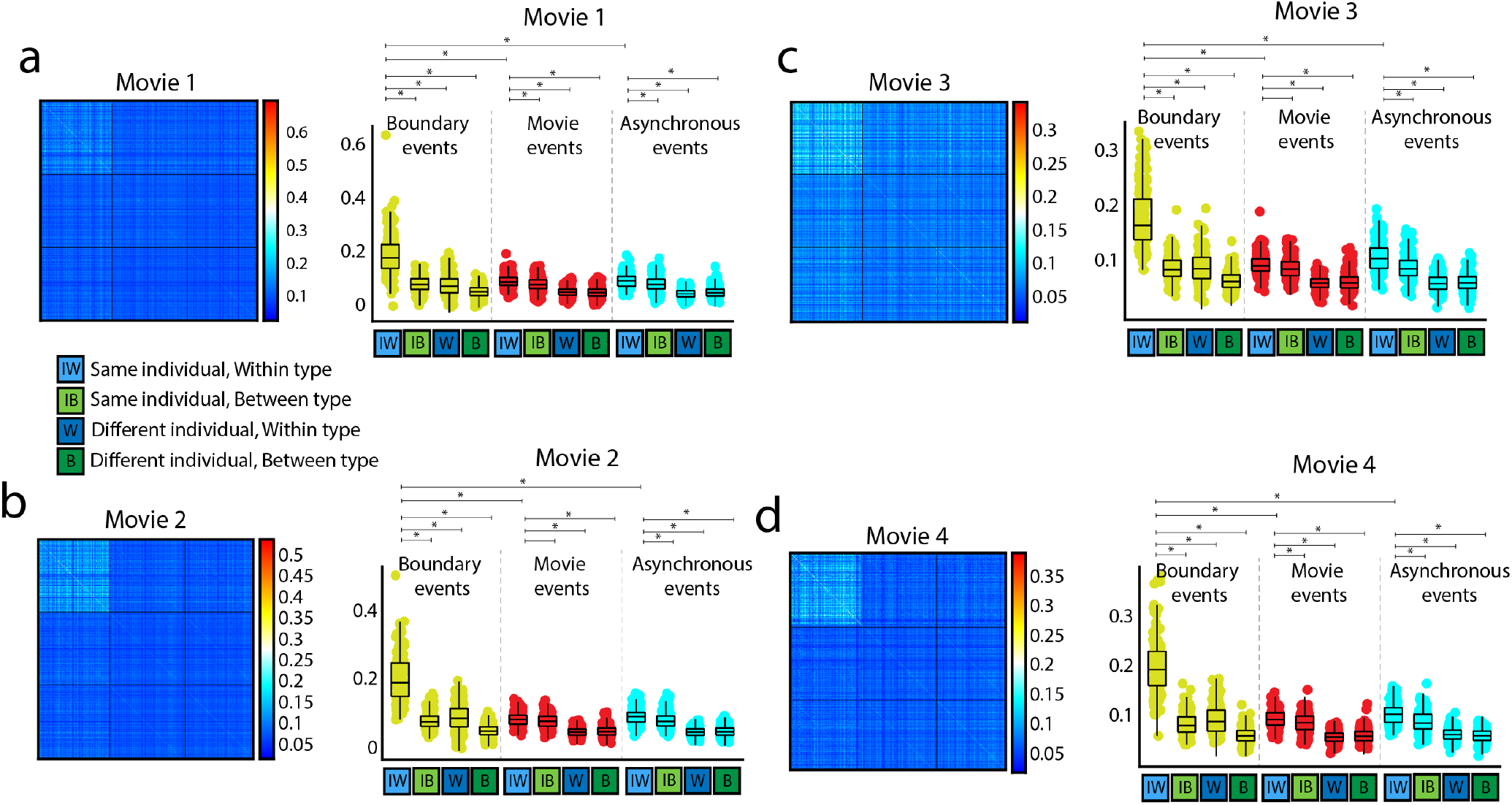
Identifiability of the event types when considered frame by frame. (*a-d*) Each of the following figures show two plots: a similarity matrix and boxplots of these values divided into four categories per event type (same individual-within type, same individual-between type, different individual-within type, and different individual-between type). All p-values below *p <* 8.66 *×* 10^*−*4^, p-values for boundary events *p <* 10^*−*15^.

**FIG. S6.**
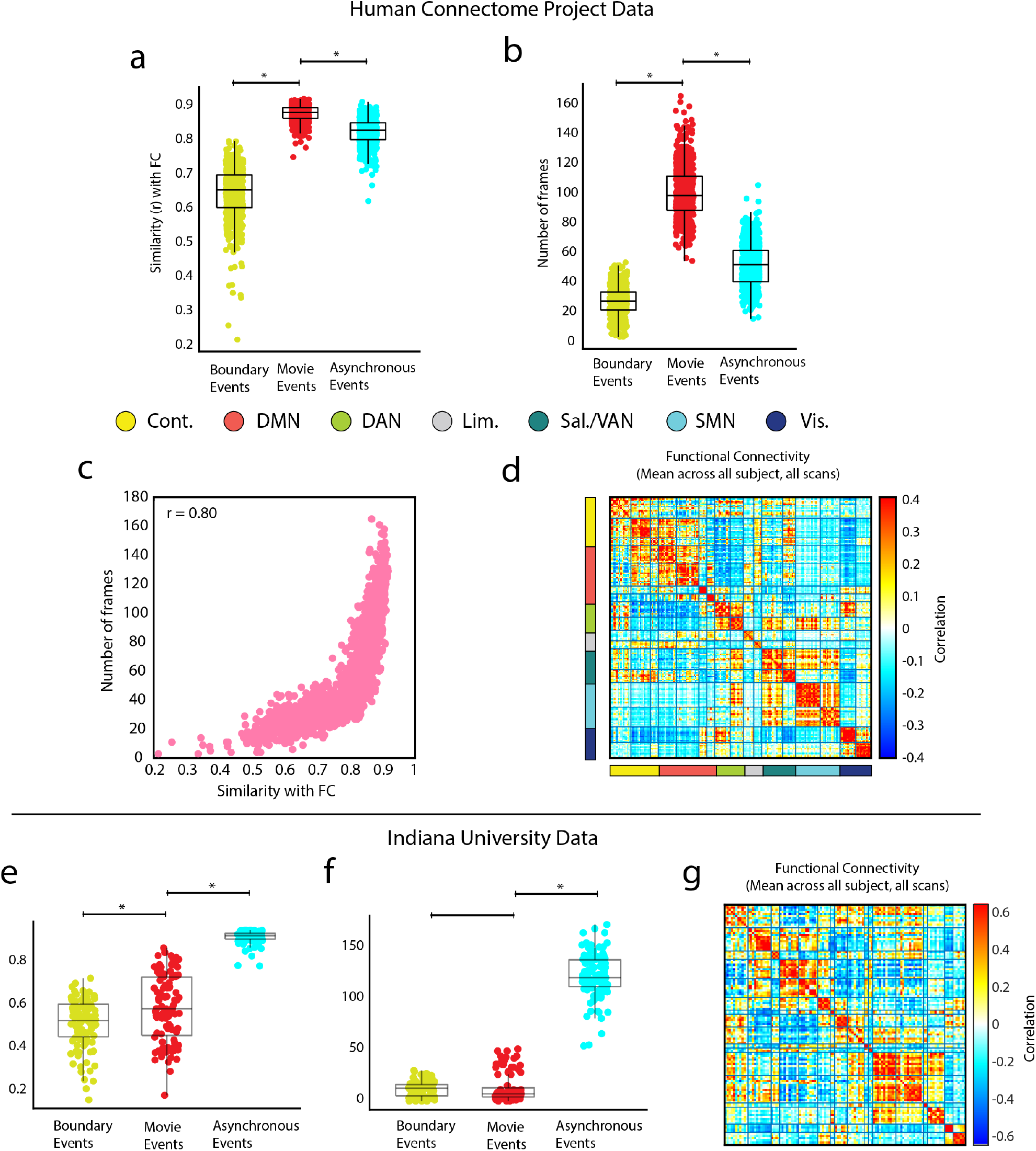
Movie events are the most similar to functional connectivity in HCP data. (*a*) Boxplots showing the distribution of correlations between an event type (mean across the participants scan) and FC for the same scan. Movie events are the most similar to FC (all p-values below *p <* 10^*−*15^). (*b*) Boxplots showing the number of frames assigned to each event type per scan/participant (all p-values below *p <* 10^*−*15^). (*c*) Scatter plot showing the relationship between similarity with FC and number of frames. Similarity with FC appears to be at least partially driven by the number of frames used to compute the mean cofluctuation pattern for an event type. (*d*) Mean functional connectivity across all scans/participants. (*e*) Boxplots showing the distribution of correlations between an event type (mean across the participants scan) and FC for the same scan. (*f*) Boxplots showing the number of frames assigned to each event type per scan/participant. (*g*) Mean functional connectivity across all scans/participants.

**FIG. S7.**
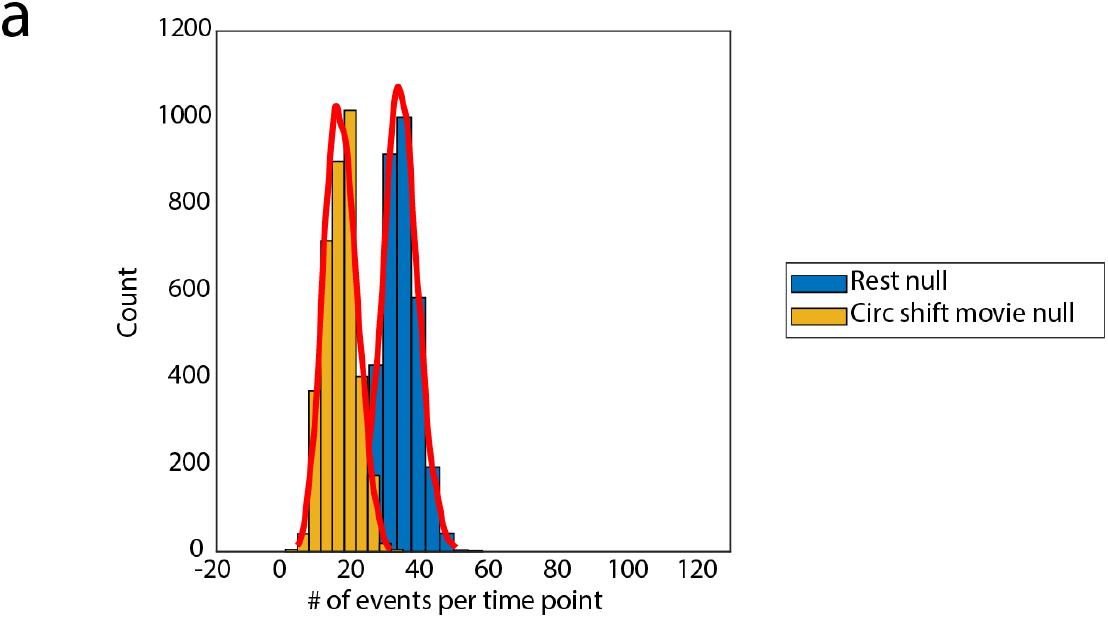
Rest-based null model is more conservative than circularly shifted movie event time series. (*a*) Plot of two null distributions. The rest null was computed as the number of events per time point while subjects were at rest. The circ shift movie null was computed as the number of events per time point with the event time series circularly shifted to maintain the number and relative timing of events in the actual data, but break the alignment.

**FIG. S8.**
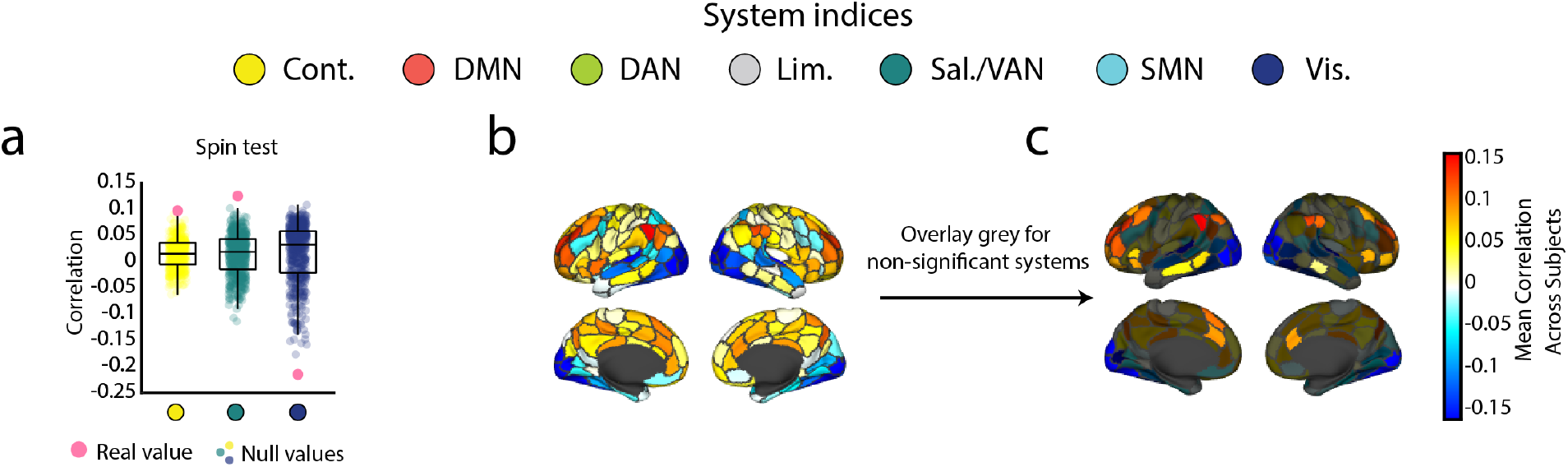
Spin test shows that correlations are significantly concentrated in control b, salience b, and central visual systems. (*a*) We found that in three systems (control b, salience b, and central visual) the mean correlation value between nodal activations in that system and boundary events was significantly higher than a null distribution created using a space-preserving permutations test (spin test). (*b*) All correlations plotted to the brains surface. (*c*) Systems that were not significant were overlaid with grey.

